# YAP nuclear translocation through dynein and acetylated microtubule controls fibroblast activation

**DOI:** 10.1101/693168

**Authors:** Eunae You, Panseon Ko, Jangho Jeong, Seula Keum, Jung-Woong Kim, Young-Jin Seo, Woo Keun Song, Sangmyung Rhee

## Abstract

Myofibroblasts are the major cell type that are responsible for increase the mechanical stiffness in fibrotic tissues. It has well documented that the TGF-β/Smad axis is required for myofibroblast differentiation under the rigid substrate condition. However, the mechanism driving myofibroblast differentiation in soft substrates remains unknown. In this research, we demonstrated that interaction of yes-associated protein (YAP) and acetylated microtubule via dynein, a microtubule motor protein drives nuclear localization of YAP in soft matrix, which in turn increased TGF-β1 induced transcriptional activity of Smad for myofibroblast differentiation. Pharmacological and genetical disruption of dynein impaired the nuclear translocation of YAP and decreased the TGF-β1 induced Smad activity even though phosphorylation and nuclear localization of Smad occurred normally in α-tubulin acetyltransferase (α-TAT1) knockout cell. Moreover, microtubule acetylation prominently appeared in the fibroblast-like cells nearby the blood vessel in the fibrotic liver induced by CCl_4_ administration which were conversely decreased by TGF-β receptor inhibitor. As a result, quantitative inhibition of microtubule acetylation may be suggested as a new target for overcome the fibrotic diseases.

## Introduction

Differentiation and phenotypical plasticity of cells are critically dependent on the nature of the cellular microenvironment (Engler, Sen et al., 2006, Kim, You et al., 2017). The cellular microenvironment is composed of various types of extracellular matrix (ECM) proteins, soluble growth factors, and numerous adjacent cell types (Joyce, 2005, Zou, 2005). The ECM proteins, which surround cells, play particularly important roles as mechanical supporters of the cells. Depending on the nature of the tissue, ECM proteins exhibit different mechanical properties with respect to the cells that constitute the tissue (Engler et al., 2006, Provenzano, Inman et al., 2009). The ability of cells to sense the mechanical properties of ECM is essential for maintaining tissue homeostasis. Disruption of tissue homeostasis by pathological processes, including tissue injury, inflammation, and cancer, is accompanied by mechanical degeneration such as fibrosis (Boutet, De Frutos et al., 2006). In this environment, the mechanically stiffened tissue allows cells to be more active, resulting in acceleration of pathological progression (Hwang, Byun et al., 2015, Jeong, Keum et al., 2018).

Fibroblasts maintain the mechanical properties of ECM by regulating the tensional status of tissues (Rhee & Grinnell, 2007). When a tissue is under pathological conditions, pro-fibrotic agonists, such as transforming growth factor-beta1 (TGF-β1), are secreted from the surrounding immune cells to promote the differentiation of fibroblasts into myofibroblasts(Wang, Qin et al., 2017). The differentiated myofibroblasts actively participate in innate stromal remodelling by changing the composition and mechanical properties of ECM during pathological progression. In fact, the number of myofibroblast expressing the alpha-smooth actin (α-SMA) as a contractile apparatus has increased in the pathological tissues including idiopathic pulmonary fibrosis (IPF), liver fibrosis, cardiovascular fibrosis and systemic sclerosis (Hardie, Glasser et al., 2009, Harris, Kelly et al., 2013). Consequently, in fibrotic tissue, an increase in tissue rigidity by myofibroblasts is the main cause of organ failure and death. Thus, understanding the mechanism of myofibroblast differentiation is important in the study of disease progression (Hinz, 2007, Tomasek, Gabbiani et al., 2002). Numerous studies have shown that ECM stiffness, in addition to biochemical agonists, is required for myofibroblast differentiation (Calvo, Ege et al., 2013). Normal lung fibroblasts cultured in stiff substrates (∼20 kPa; lung fibrotic rigidity) without agonist, acquire a myofibroblastic phenotype including α-SMA expression than (∼0.5 kPa; normal lung rigidity), which is induced by integrin-mediated mechanotransduction pathway (Gabbiani, 2003, Hinz, Phan et al., 2007, Huang, Yang et al., 2012, You, Huh et al., 2019). However, because early pathological tissues are as soft as normal tissues, the signalling pathways associated with myofibroblast differentiation in soft substrates may differ from those occurring in stiff substrates. Nevertheless, the molecular mechanisms driving myofibroblast differentiation in soft substrates remain unclear. We have also recently found that mouse embryonic fibroblasts (MEFs) with knockout (KO) of the *Spin90* gene show increased microtubule acetylation and myofibroblastic phenotype in a 0.5 kPa polyacrylamide gel (PAG) substrate that mimics normal tissue stiffness, indicating that microtubules may be involved in myofibroblast differentiation under soft substrate conditions (You, Huh et al., 2017 b57). However, whether microtubule dynamics are required for myofibroblast differentiation under pathological conditions remains to be elucidated.

Yes-associated protein (YAP) is a key regulator of myofibroblast response to ECM stiffness. Notably, inhibition of integrin-derived cellular contractile activity via inhibition of myosin and Rho kinase activities significantly reduces YAP nuclear accumulation, indicating that YAP is activated by intrinsic mechanotransduction in response to ECM stiffening (Maller, DuFort et al., 2013 b36). Altogether, YAP/TAZ complex mediates the mechanical stress exerted by the ECM and drives a positive-feedback loop to accelerate fibrotic disease and cancer development. Interestingly, cytoplasmic YAP under low matrix stiffness is translocated into the nucleus upon treatment of TGF-β1 in dermal fibroblasts, suggesting the existence of a mechanism in which YAP is translocated to the nucleus when cells are grown on a soft matrix (Piersma, de Rond et al., 2015). However, the underlying molecular mechanisms responsible for TGF-β1-induced nuclear entry of YAP under low matrix stiffness remain unknown.

Cellular signalling differs in 2D vs. 3D collagen matrices (Rhee & Grinnell, 2007, Rhee, Jiang et al., 2007), which impacts myofibroblast differentiation. In addition, microtubule acetylation is required for the myofibroblast differentiation under soft substrate condition. These findings highlight the need to understand how microtubule acetylation controls the myofibroblast differentiation under soft substrate condition. In this study, we investigated the molecular mechanisms driving myofibroblast differentiation in soft matrices that mechanically mimic the early pathological stage of fibrosis.

## Results

### Myofibroblast differentiation induced by TGF-β1 is accompanied by microtubule acetylation in cells grown on soft matrices

To compare the effects of TGF-β1 on myofibroblast differentiation with respect to substrate stiffness, MEFs were seeded on 0.5 kPa PAG substrates (soft) and glass coverslips (stiff) and cultured for 8 h with or without the presence of TGF-β1. As shown in Fig. 1a, TGF-β1 induced strong focal adhesions and formation of actin stress fibres in cells cultured on glass coverslips, but not in cells cultured on soft matrices. Formation of strong focal adhesions and actin stress fibres was nearly absent in cells cultured on soft PAG matrices. Instead, the prominent features of cells cultured on soft PAG matrices included increased cell length, and increased ratio of nuclear length to width (represented by the nuclear elliptical factor [EF]) after treatment with TGF-β1 (Fig 1A). Consistent with the results of previous studies, the spreading of fibroblasts on soft matrices (Rhee et al., 2007), but not on stiff matrices, was completely inhibited by treatment with nocodazole, a microtubule disrupting agent (Fig 1B). This result indicates that microtubules play an important role in the spreading of cells on soft matrices under TGF-β1 stimulation.

**Figure 1.**
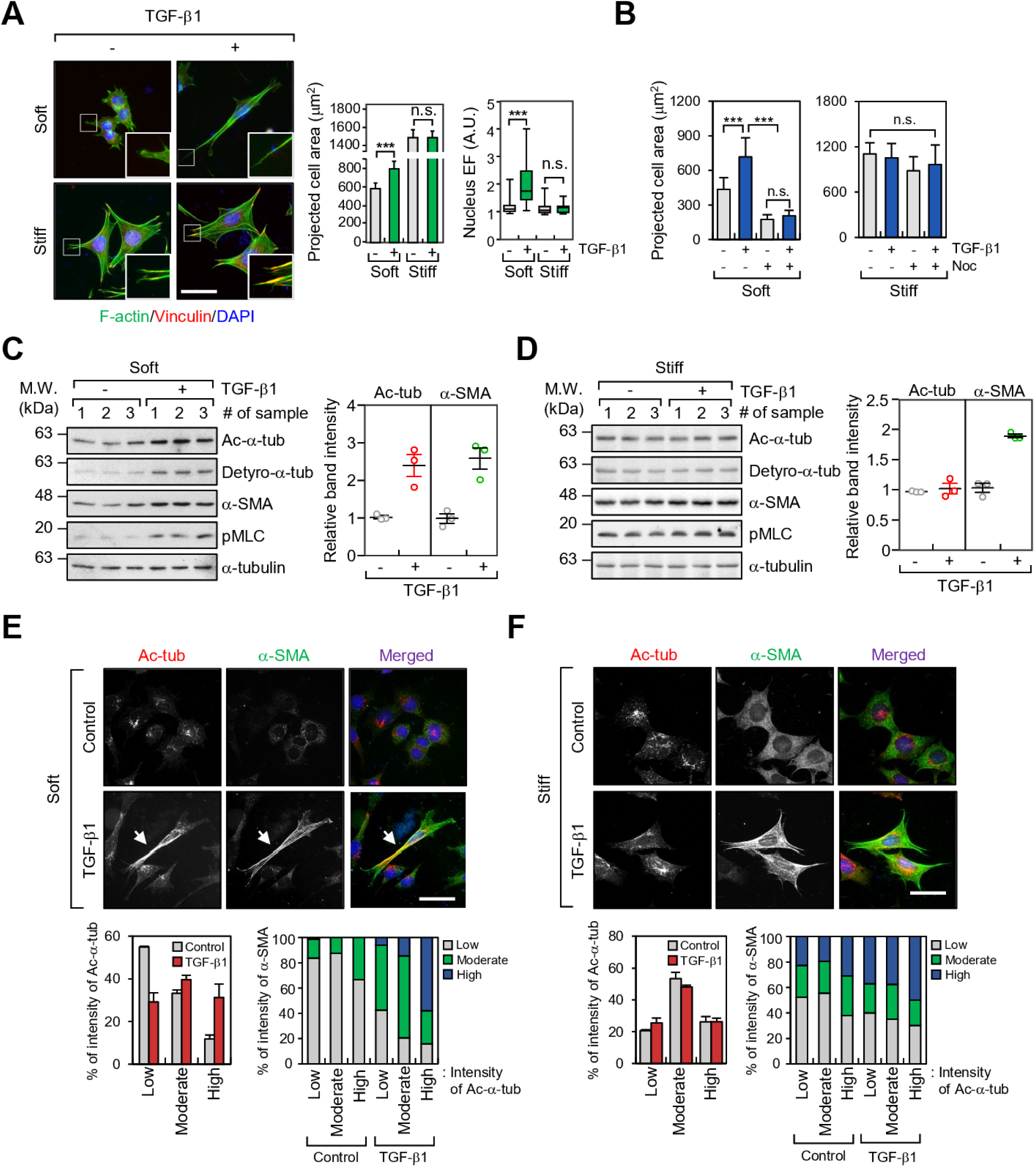
TGF-β1 increases microtubule acetylation on soft matrices. A Mouse embryonic fibroblasts (MEFs) were seeded on soft (0.5 kPa PAGs) and stiff (glass) conditions, stimulated with TGF-β1 (2 ng ml^-1^) for 8 h, and then labelled using an antibody specific for vinculin and Alexa Fluor^®^ 488 phalloidin. Graphs show quantification of projected cell area and nuclear elliptical factor (EF) for at least 30 cells under each condition. Error bars represent S.D. of the data. \*\*\**p* < 0.005; n.s. indicates non-significant. Scale bar, 50 μm. B Comparison of projected cell area upon combinational treatment using TGF-β1 and/or nocodazole (Noc; 10 μM). \*\*\**p* < 0.001; n.s. indicates non-significant. C, D Western blotting conducted to detect the level of post-translational modifications including acetylation and detyrosination, α-tubulin and α-SMA in TGF-β1-treated MEFs incubated on soft (C) and stiff (D) matrices. Protein lysates, obtained from MEFs grown on soft and stiff, were prepared independently. Graph shows relative expression of α-tubulin acetylation (upper) and α-SMA (lower) normalized to the expression of α-tubulin. E, F Immunofluorescence labelling of acetylated-α-tubulin and α-SMA in MEFs incubated on soft (E) and stiff (F) matrices upon stimulation with TGF-β1. Expression of acetylated-α-tubulin and α-SMA was categorized using fluorescence intensity (low, moderate, and high). Graphs show the percentage of α-SMA expression as increasing fluorescence intensity of acetylated-α-tubulin. Arrows indicate co-expression of acetylated-α-tubulin and α-SMA in TGF-β1-treated MEFs grown under soft conditions. Scale bar, 50 μm.

Previously, we reported that microtubule acetylation is critical for myofibroblast differentiation in SPIN90-depleted MEFs in a soft matrix environment (You et al., 2017). In the present study, we confirmed that microtubule acetylation is also increased during myofibroblast differentiation induced by TGF-β1. Interestingly, acetylation and detyrosination of microtubule were significantly increased during myofibroblast differentiation induced by TGF-β1 on soft matrices, which was due to increased α-SMA expression (Fig 1C; soft matrix). In contrast, those modifications did not induce by TGF-β under stiff substrate condition (Fig 1D; stiff matrix). In addition to TGF-β1, LPA, another inducer of myofibroblast differentiation (Mazzocca, Dituri et al., 2011), increased the acetylation of microtubules and expression of α-SMA under soft substrate conditions (Figure EV1A and B). Fibroblasts cultured on soft matrices show reduced integrin-mediated focal adhesion signalling (Mih, Marinkovic et al., 2012, Rhee et al., 2007). Therefore, we hypothesized that increased microtubule acetylation in fibroblasts cultured on a soft substrate is associated with absence of integrin-mediated signalling. To test this hypothesis, we examined the extent of TGF-β1-induced microtubule acetylation after treatment with pharmacological inhibitors of integrin signalling in cells cultured on stiff. Fibroblasts treated with blebbistatin (myosin II inhibitor) and Y27632 (Rho-associated kinase inhibitor) showed an increase in microtubule acetylation induced by TGF-β1 even when cultured on a stiff substrate (Figure EV2A and B). This result indicates that acetylated microtubules play a prominent role in myofibroblast differentiation under conditions of weak integrin signalling, such as when cultured on soft matrices and in a 3D environment.

To investigate the correlation between microtubule acetylation and expression of α-SMA, we examined its cellular localization in TGF-β1-treated MEFs cultured on soft and stiff matrices. The fluorescence intensities of α-SMA and acetylated-α-tubulin were arbitrarily divided into three categories; high, moderate, and low. Fibroblasts cultured on soft matrices exhibited high expression of α-SMA as the levels of acetylated-α-tubulin increased after stimulation with TGF-β1 (Fig 1E). However, the relationship between acetylated-α-tubulin and α-SMA in fibroblasts cultured under stiff substrate conditions was unclear (Fig 1F). We also found acetylated microtubules in cells cultured on soft substrates were distributed along the length of the cell, whereas in cells cultured on stiff substrates, they were present around the nucleus (Fig 1E and F). These findings indicate that acetylated microtubules in the cell cultured under soft matrices are critical for cell spreading and morphogenesis.

### Microtubule acetylation is required for TGF-β1-induced myofibroblast differentiation on soft matrices

To analyze the molecular mechanisms of microtubule acetylation in myofibroblast differentiation on soft substrates, we generated an MEF cell line in which the α-TAT1, encoded by *Atat1* gene was disrupted using the CRISPR/Cas9 system (α-TAT1 KO MEF; Fig 2A, Fig Figure EV3A and B). We were unable to directly confirm the ablation of endogenous α-TAT1 protein expression in α-TAT1 KO MEFs via western blotting because there are no commercially available antibodies that bind the endogenous α-TAT1. However, two clones of the α-TAT1 KO MEF cell line were selected using DNA sequencing of the CRISPR/Cas9 target site in the *Atat1* gene and by the yield of tubulin acetylation (Fig 2A). α-TAT1 KO MEFs showed a profound decrease in TGF-β-induced nuclear elongation (Fig 2B). α-SMA (encoded by *Acta2* gene) transcription and protein expression were also inhibited in α-TAT1 KO MEFs cultured on soft substrates, while the level of detyrosinated tubulin, assessed under the same conditions, remained unchanged (Fig 2C). Cells incubated on a stiff substrate increased their expression of α-SMA in response to stimulation with TGF-β1 regardless of the presence of acetylated-α-tubulin (Fig 2D). Fluorescence imaging confirmed that the expression of α-SMA induced by TGF-β occurred in α-TAT1 KO MEF, but was less than that in WT MEFs (Fig 2E). Transient overexpression of GFP-α-TAT1 in WT MEFs increased the expression of α-SMA in response to stimulation with TGF-β (Fig 2F), suggesting that acetylated microtubules are involved in TGF-β1-mediated myofibroblast differentiation on soft substrates.

**Figure 2.**
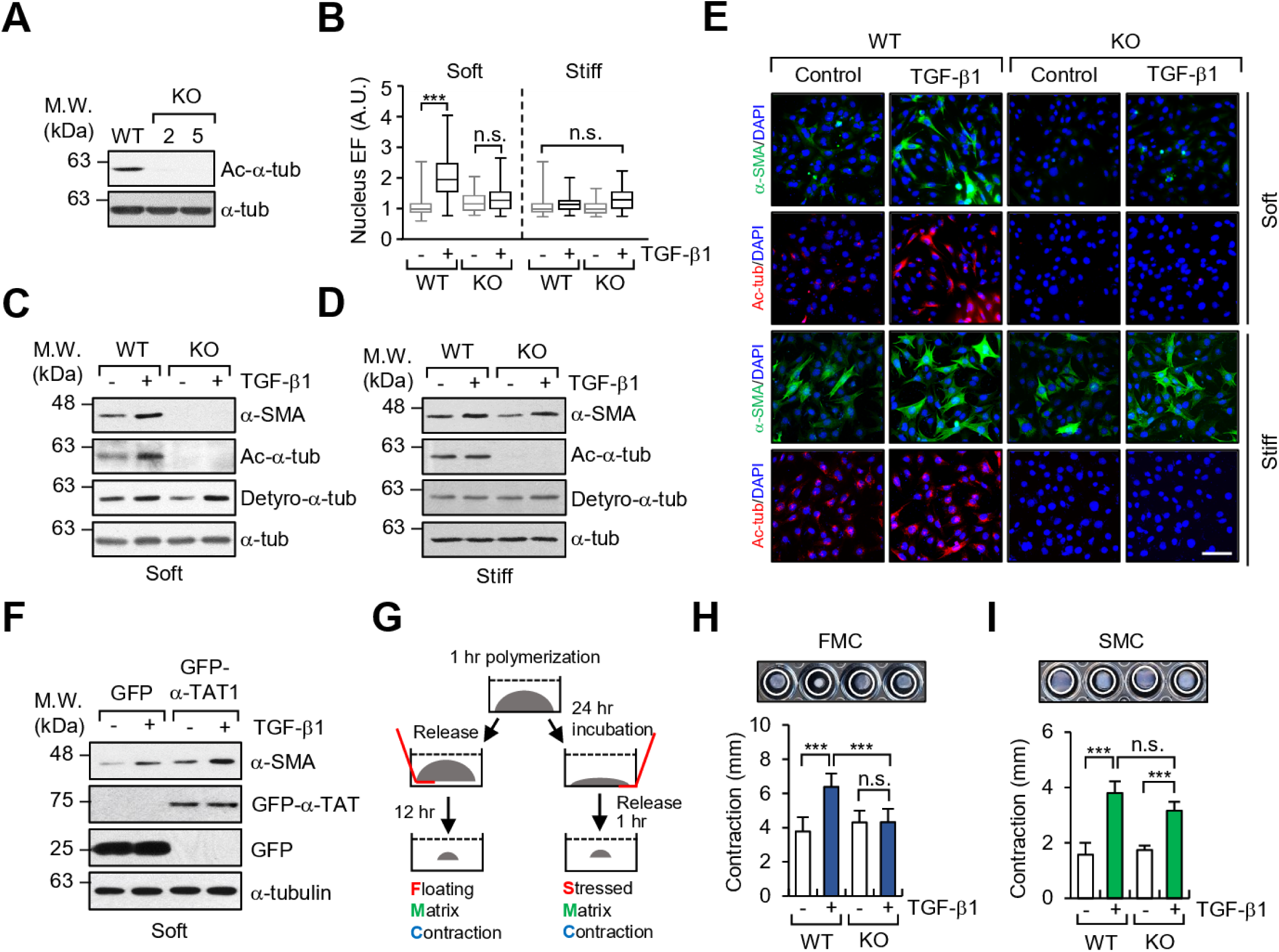
Increase in microtubule acetylation is required for TGF-β1-induced myofibroblast differentiation on soft matrix. A Western blotting conducted to detect the level of acetylated-α-tubulin in α-TAT1 KO MEFs (clone #2 and 5) established using a CRISPR/Cas9 system. B Quantification of nuclear elliptical factor (EF) in WT and α-TAT1 KO MEFs upon treatment with TGF-β1 for 8 h. \*\*\**p* < 0.005; n.s. indicates non-significant. C, D Western blotting of α-SMA in WT and α-TAT1 KO MEFs grown on soft (C) and stiff (D) matrices. E Immunofluorescence images of α-SMA (green) and acetylated-α-tubulin (red) in WT and α-TAT1 KO MEFs upon stimulation with TGF-β1. Scale bar, 100 μm. F MEFs were transfected with GFP or GFP-α-TAT1 for 24 h. After serum starvation for 12 h, cells were seeded on fibronectin-coated soft matrix and stimulated with TGF-β1 for 8 h. Cells were lysed and subjected to western blotting using antibodies specific for α-SMA and GFP. G Schematic diagram of 3D collagen matrix remodelling assays (FMC, floating matrix contraction; SMC, stressed matrix contraction). H, I WT and KO cells were embedded in 3D matrices composed of 1 mg ml^-1^ collagen and 100 μg ml^-1^ fibronectin, and incubated for 1 h. After adding TGF-β1 into each matrix, FMC (H) and SMC (I) assays were performed as shown in (G). Graphs show the reduced size of the 3D collagen gel compared with original gel size. Data represent the mean of three independent experiments ± S.E.M. \*\*\**p* < 0.005.

We next investigated whether acetylated microtubules are required for fibroblast contractility and re-organization of 3D collagen matrix (Fig 2G). Fibroblasts in floating matrix contraction (FMC) initially have a round morphology and then spread during contraction, which resembles cells in soft matrices. Fibroblasts in stressed matrix contraction (SMC) show a spread morphology with stress fibres, and resemble the fibroblasts cultured on stiff substrates (Grinnell, Ho et al., 1999, Rhee & Grinnell, 2007). As shown in Fig 2H and I, α-TAT1 KO MEFs, stimulated with TGF-β1, did not increase the FMC, whereas both WT and α-TAT1 KO MEFs induced SMC to a similar extent in response to stimulation with TGF-β. This result indicates that acetylated microtubules play an important role in contractility of cells grown on soft substrates before substrate develops tension.

### Microtubule acetylation is involved in TGF-β1-induced expression of myofibroblast marker genes in MEFs grown on a soft substrate

To examine whether microtubule acetylation induces gene expression associated with myofibroblast differentiation in fibroblasts grown on a soft matrix, we performed RNA-sequencing analysis to compare the differentially expressed genes (DEGs) between WT and α-TAT1 KO MEFs seeded on a soft substrate in the absence or presence of TGF-β1 (Fig 3A). The expression of 657 genes was significantly increased and that of 574 genes was significantly downregulated upon TGF-β1 stimulation (±1.4 fold change, *p*-value < 0.05; Fig 3B, left). Among the 657 genes that were upregulated in WT MEFs upon stimulation with TGF-β1, 83 were downregulated in α-TAT1 KO MEFs (−1.4 fold change, *p*-value < 0.05) (Fig 3B, right). The biological process regulated by these 83 genes were assessed by gene ontology (GO) analysis (Fig 3D). Of these 83 genes, 32 belonged to the top tier of genes for cellular process; the majority of these genes was involved in metabolic and cellular localization processes (Fig 3C; left panel). We further investigated the specific functions of these genes involved in cellular processes, and found that genes regulating organization of cellular components, such as that of cytoskeletal binding proteins, appeared the highest level (Fig 3C; right panel). During myofibroblast differentiation, cytoskeletal proteins play crucial roles in cell migration and contraction (Cai, Chou et al., 2012, Gimona, Sparrow et al., 1992, Rockey, Weymouth et al., 2013); thus, genes whose expression is regulated by α-TAT1 are likely involved in cytoskeletal protein rearrangement and signal transduction during myofibroblast differentiation.

**Figure 3.**
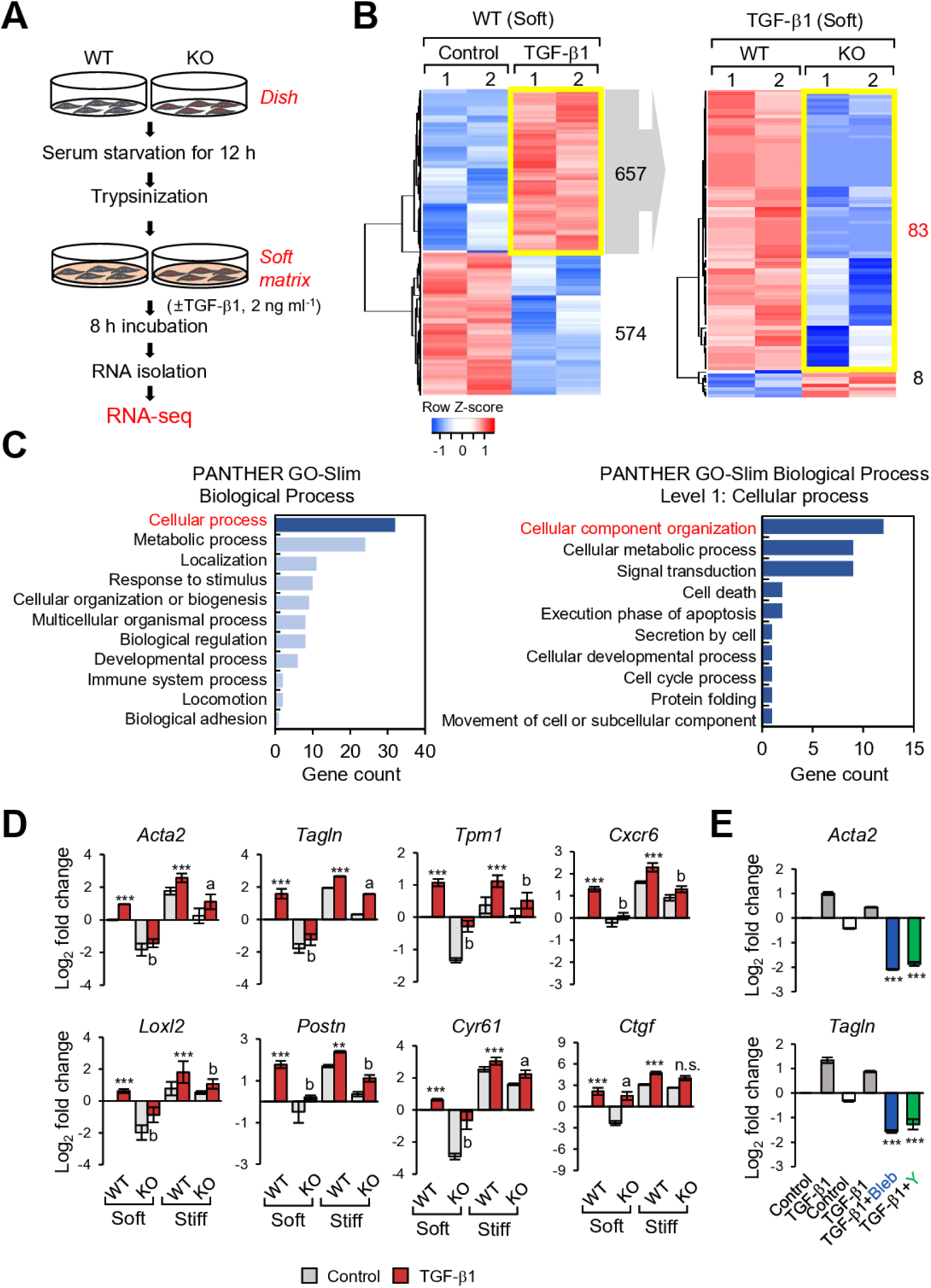
Differential gene expression in WT and α-TAT1 KO MEFs upon stimulation with TGF-β1. A Schematic diagram of the RNA-seq analysis workflow. B Left; Heatmap shows differentially expressed genes (DEGs; upregulated: 657; down-regulated: 574) in WT MEFs stimulated with TGF-β1. Right; DEGs between WT and KO upon TGF-β1 stimulation among 657 genes upregulated by TGF-β1. C Functional annotation of 83 genes selected on (B; right) by Panther gene ontology. D Validation of myofibroblast marker genes using RT-qPCR. Data represent the mean of three independent experiments ± S.E.M. \**p* < 0.05, \*\**p* < 0.01, \*\*\**p* < 0.005 as compared with WT (control). ^a^*p* < 0.05, ^b^*p* < 0.005 as compared with WT stimulated with TGF-β1. E Transcript levels of *Acta2* and *Tagln* in α-TAT1 KO MEFs after combinational treatment with TGF-β1 and blebbistatin (Bleb)/or Y27632 (Y), incubated under stiff conditions. Data represent the mean of three independent experiments ± S.E.M. \*\*\**p* < 0.005.

Next, we determined the expression levels of 6 genes (*Acta2, Tagln, Tpm1, Cxcr6, Postn*, and *Loxl2*) out of 83 genes associated with myofibroblast differentiation that were identified as upregulated genes in the WT cells treated with TGF-β1 compared with KO. In addition, because *Ctgf* and *Cyr61* genes were identified as significantly upregulated genes depending on microtubule acetylation (You et al., 2017), we also compared these genes expression by RT-qPCR analysis in soft and stiff conditions (Fig 3D). RT-qPCR performed with samples obtained using cells cultured on soft matrices, showed that the 8 examined genes were upregulated by treatment with TGF-β1, whereas in α-TAT1 KO MEFs, treatment with TGF-β1 failed to induce the expression of these genes. Notably, differences in the gene expression induced by TGF-β1 in WT and α-TAT1 KO were not remarkable under stiff matrix conditions (Fig 3D). In addition, treatment with blebbistatin and Y27632 dramatically inhibited the expression of *Acta2* and *Tagln*, decreasing it to basal levels in α-TAT1 KO MEFs under stiff matrix conditions (Fig 3E). These results suggest that acetylated microtubules are indispensable for gene expression associated with myofibroblast differentiation in fibroblasts grown on a soft substrate.

### Microtubule acetylation are required for YAP-and Smad-dependent transcriptional activity

We have previously reported that microtubule acetylation is associated with nuclear translocation of the YAP protein and induction of myofibroblast differentiation in MEFs grown on soft matrices (You et al., 2017). In addition to YAP, the Smad transcription factor is also involved in the TGF-β-induced myofibroblast differentiation (de Caestecker, Parks et al., 1998, Nakao, Imamura et al., 1997). Thus, we explored whether acetylated microtubules also regulate Smad activity in TGF-β-mediated myofibroblast differentiation under soft matrix conditions. MEF cell lines with a knockdown (KD) of YAP and Smad2 were established using shYAP and sh*S*mad2 viral vector expression. The expression of Smad and YAP in KD cells was knocked down by ∼70% in transcript and protein levels (Fig 4A). Analysis of myofibroblast marker gene expression upregulated by acetylation of microtubules, showed that both Smad2 and YAP KD MEFs were inhibited to a similar extent in the expression of all tested genes, however, such as *Cyr61* and *Ctgf* in Smad2 KD, and *Postn* and *Cxcr6* in YAP KD did not show inhibited expression (Fig 4B). Matrix contractility, mediated by TGF-β1, was also significantly inhibited by the knockdown of Smad2 and YAP, respectively (Fig 4C), indicating that contractile activity in Smad2 or YAP KD fibroblasts was similar to that in α-TAT1 KO fibroblasts.

**Figure 4.**
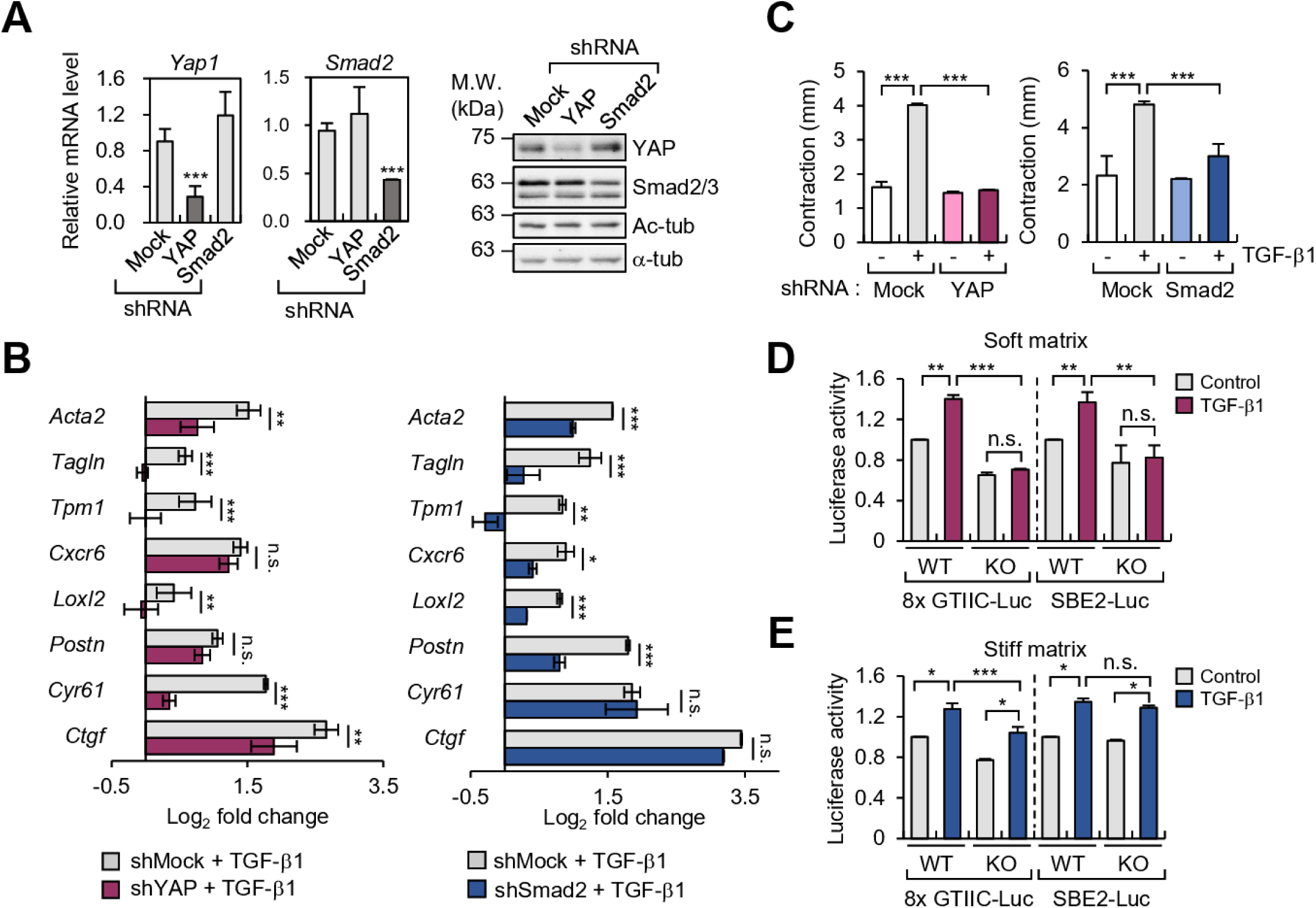
Increased acetylation of microtubules, induced by TGF-β1, regulates myofibroblast differentiation in YAP/Smad2-dependent manner. A Transcripts and protein levels of YAP and Smad2 in MEFs expressing pLKO.1-bla-shSmad2 and pLKO.1-bla-shYAP1. Data represent the mean of three independent experiments ± S.E.M. \*\*\**p* < 0.005. B RT-qPCR analysis conducted to compare gene expression in MEFs expressing pLKO.1-bla-shSmad2 and pLKO.1-bla-shYAP1. Relative transcript level was normalized using pLKO.1-bla-transfected cells. Data represent the mean of three independent experiments ± S.E.M. \**p* < 0.05, \*\**p* < 0.01, \*\*\**p* < 0.005. C FMC assay conducted to evaluate the activity of pLKO.1-bla-shSmad2 and pLKO.1-bla-shYAP1 upon stimulation with TGF-β1. Graphs indicate reduced diameter of a 3D collagen gel compared with that of the original size. \*\*\**p* < 0.005. D, E Luciferase reporter assay conducted to measure the transcriptional activity of YAP (8xGTIIC-Luc) and Smad (SBE2-Luc). WT and KO MEFs were transfected using SBE2-Luc or 8xGTIIC-Luc with CMV-β-galactosidase for 24 h. Cells were serum starved for 12 h, seeded on fibronectin-coated soft (D) or stiff matrices (E), and incubated for 8 h with TGF-β1. Luciferase activities were normalized using the activity of β-galactosidase. Data represent the mean of three independent experiments ± S.E.M. \**p* < 0.05, \*\**p* < 0.01, \*\*\**p* < 0.005; n.s. indicates non-significant.

To determine whether acetylated microtubules control the transcriptional activity of YAP and Smad in MEFs grown on a soft substrate, we performed a reporter assay for Smad and YAP using WT and α-TAT1 KO MEFs under soft and stiff matrix conditions (Fig 4D and E). Interestingly, the transcriptional activity of Smad and YAP were dramatically reduced in α-TAT1 KO MEFs compared with that of WT MEFs stimulated with TGF-β1 under soft substrate conditions (Fig 4D). Together, these findings indicate that acetylated microtubules are key regulators of TGF-β1-mediated YAP and Smad transcriptional activity in fibroblasts under soft substrate conditions but not under stiff substrate conditions.

### YAP entered the nucleus via acetylated microtubule and increases Smad activity in soft substrate

The abovementioned results indicate that acetylated microtubules are required for YAP and Smad activities in soft substrate. Therefore, we next examined whether nuclear localization of phospho-Smad2/3 and YAP in cells grown on a soft substrate and stimulated with TGF-β1 is also regulated by acetylated microtubules. As shown in Fig 5A, stimulation with TGF-β1 induced complete nuclear translocation of phospho-Smad2/3 in both WT and α-TAT1 KO MEFs. The amount of phospho-Smad2/3 in the nuclei of WT and α-TAT1 KO MEFs was similar. However, the less amount of nuclear YAP was detected in response to stimulation with TGF-β1 in α-TAT1 KO MEFs compared with those in WT MEFs under soft substrate conditions (Fig 5A and Figure EV4). Conversely, in cells grown on a stiff matrix, stimulation with TGF-β1 robustly induced the nuclear localization of phospho-Smad2/3 irrespective of the presence or absence of acetylated microtubule.

**Figure 5.**
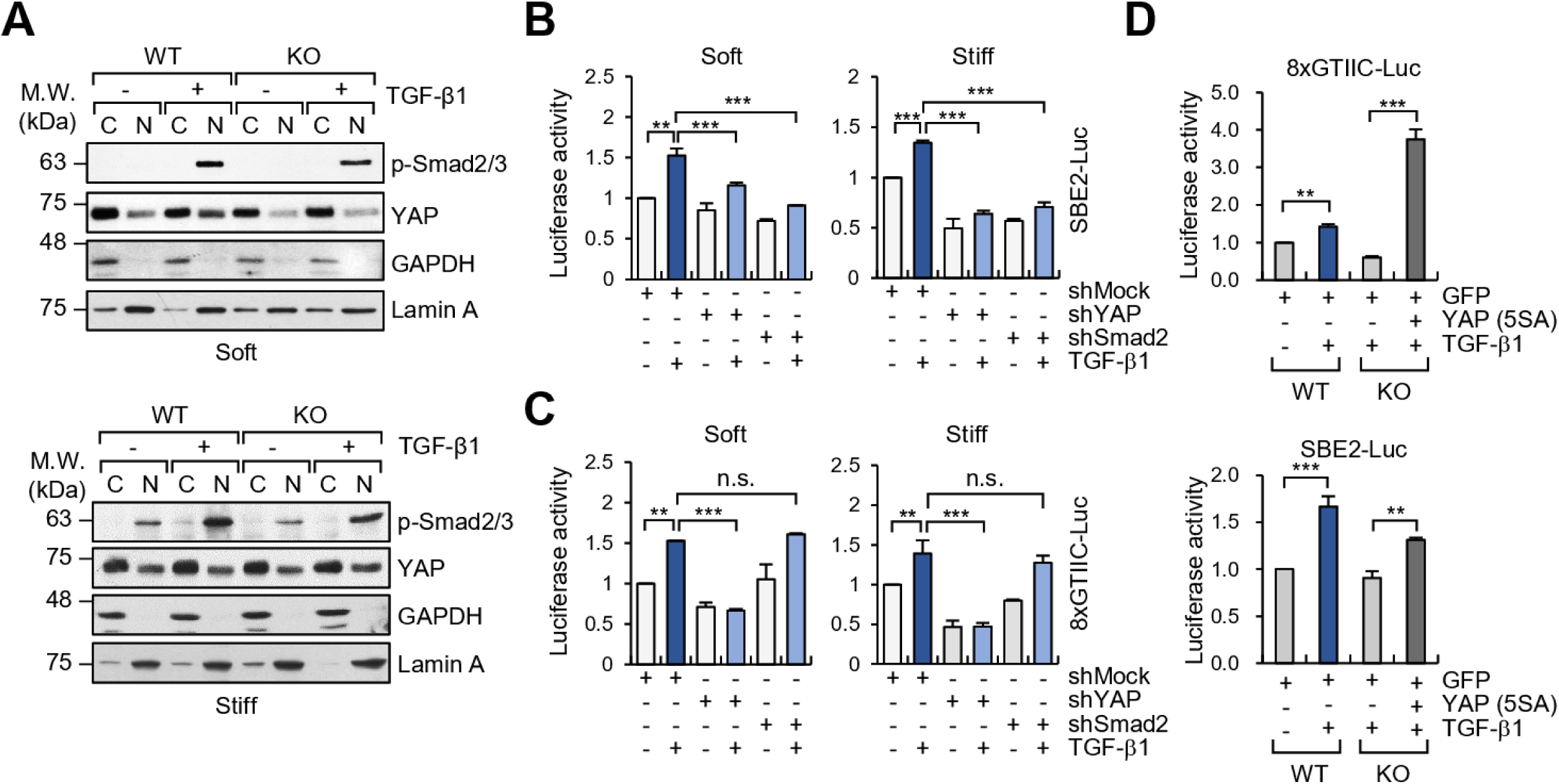
Microtubule acetylation regulates TGF-β1-induced nuclear translocation of YAP, resulting in promotion of Smad2 transcriptional activation. A Subcellular fractionation in WT and α-TAT1 KO MEFs incubated on soft and stiff matrices and stimulated with TGF-β1. Each fraction was used for western blotting with antibodies specific for phospho-Smad2/3 and YAP. GAPDH and Lamin A were used as markers for cytosolic and nucleic fraction, respectively. B, C Luciferase reporter assay for transcriptional activity of Smad (B; SBE2-Luc) and YAP (C; 8xGTIIC-Luc) in YAP and Sma2 KD cells incubated on soft or stiff matrices. \*\**p* < 0.01, \*\*\**p* < 0.005. n.s. indicates non-significant. D, E WT and α-TAT1 KO MEFs were transfected with the indicated plasmids (GFP or YAP(5SA)-GFP) for 24 h. WT and α-TAT1 KO MEFs were seeded on fibronectin-coated soft matrix and incubated for 8 h. Graphs show the relative luciferase activity of YAP and Smad. \*\**p* < 0.01, \*\*\**p* < 0.005.

Unstable phospho-Smad2/3 induce the de-phosphorylation and exported it out of the nucleus (Lin, Duan et al., 2006, Szeto, Narimatsu et al., 2016). Thus, it is possible that nuclear YAP, translocated by acetylated microtubules, can maintain the phosphorylated status of Smad2/3 in the nucleus, thereby assisting the activity of Smad2/3 in the cell grown in soft substrates. As we expected, the knockdown of YAP significantly reduced the transcriptional activity of Smad2/3 induced by TGF-β1, while knockdown of Smad did not influence the transcriptional activity of YAP/TEAD (Fig 5B and C). These findings are consistent with the results showing that inhibition of YAP with verteporfin, a pharmacological inhibitor, attenuates TGF-β1-induced nuclear accumulation of phospho-Smad, resulting in reduced expression of TGF-β1 target genes (Szeto et al., 2016). Overexpression of YAP active mutant (5SA) restored the transcriptional activity of Smad in α-TAT1 KO MEFs (Fig 5D). Taken together, these results support the hypothesis that nuclear translocation of YAP, induced by acetylated microtubules, promotes transcriptional activation of Smad2/3 in MEFs grown on soft matrices.

### TGF-β1-induced YAP is translocated along microtubule-dynein complex

Microtubule acetylation facilitates the accessibility of the motor proteins kinesin-1 and dynein to microtubules (Reed, Cai et al., 2006). Increased accessibility of these motor proteins to acetylated microtubules promotes anterograde and retrograde transport of cargo proteins. Therefore, we examined whether the nuclear translocation of YAP by acetylated microtubules in a soft matrix is dependent on the motor protein dynein. To disturb dynein activity, WT MEFs were treated with pharmacological inhibitor of dynein, erythro-9-(2-hydroxy-3-nonyl)adenine (EHNA), which blocks ATPase and motor activity (Lin & Nicastro, 2018). EHNA robustly inhibited TGF-β1-induced translocation of YAP into the nucleus without affecting cell morphology (Fig 6A and B). Overexpression of dynamitin, which is reported to inhibit cytoplasmic dynein-based motility by inducing disassembly of dynactin (Burkhardt, Echeverri et al., 1997), also dramatically inhibited TGF-β1-induced nuclear translocation of YAP on the soft substrate (Figure EV5A and B). Conversely, nuclear localization of phospho-Smad2/3 was not inhibited by EHNA, confirming the notion that nuclear translocation of phospho-Smad2/3 is independent of dynein activity (Fig 6B). To further explore whether dynein indeed interacts with YAP via acetylated microtubule, we performed a microtubule sedimentation assay. When cells were treated with TGF-β1, YAP readily precipitated with dynein in WT, but not in α-TAT1 KO MEFs, indicating that YAP can form a tertiary complex with dynein and acetylated microtubules (Fig 6C).

**Figure 6.**
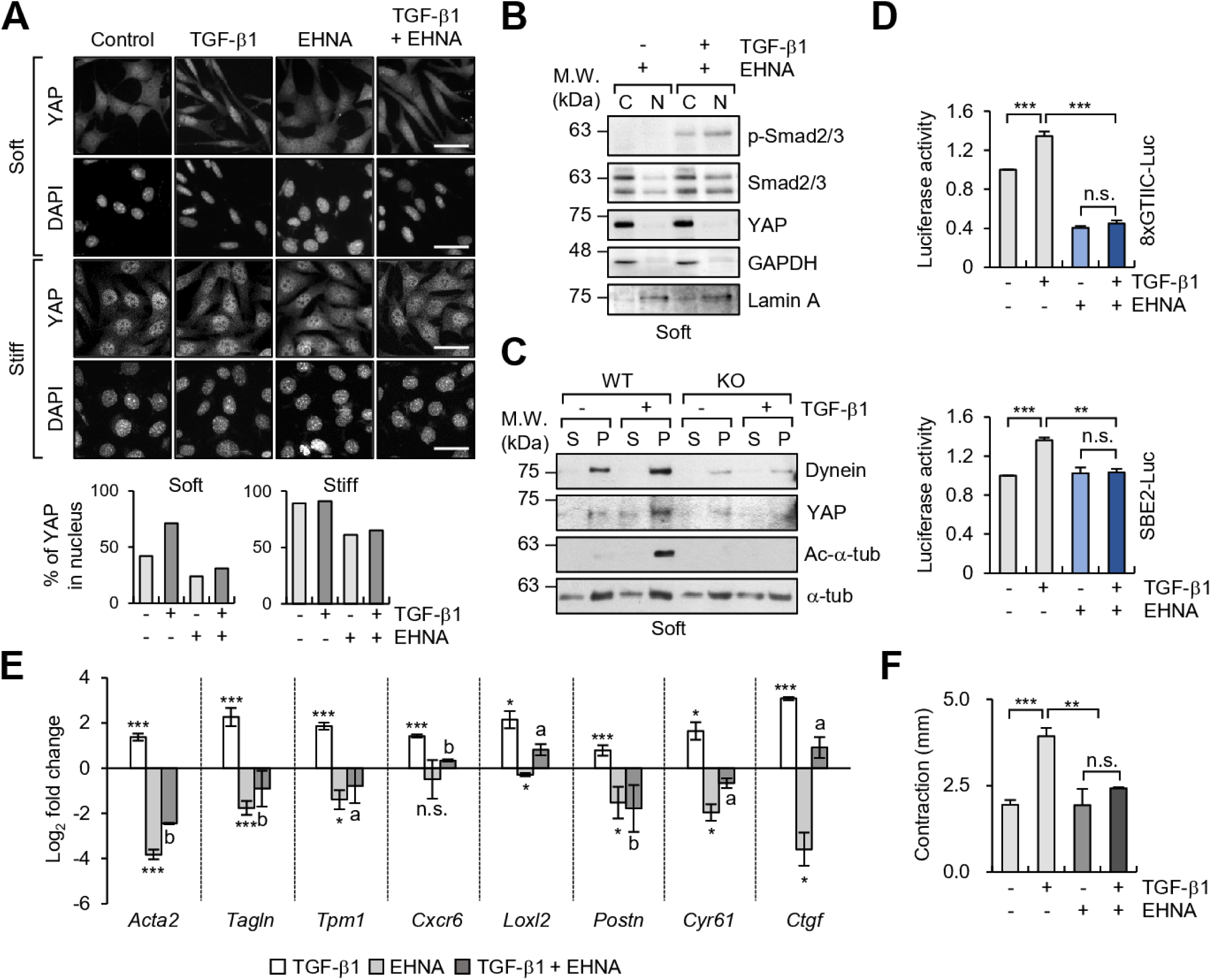
Acetylated microtubules are sufficient for dynein-dependent nuclear translocation of YAP upon stimulation with TGF-β1. A Immunolabeling of YAP expression in WT and α-TAT1 KO MEFs treated with EHNA (500 μM) and/or TGF-β1 (2 ng ml^-1^) and grown on fibronectin-coated soft and stiff matrices. Graphs show the percentage of YAP in the nucleus. Scale bar, 50 μm. B Western blotting analysis of nuclear and cytosolic fractions of WT MEFs treated with EHNA and/or TGF-β1 for 8 h. GAPDH and Lamin A were used as markers for cytosolic and nucleic fractions, respectively. C Microtubule sedimentation assay examining WT and α-TAT1 KO MEFs incubated on a soft matrix. Each fraction was assessed via western blotting to detect the levels of dynein and YAP. S; supernatant (depolymerized tubulin), P; pellet (polymerized tubulin). D Luciferase reporter assay (8XGTIIC-Luc and SBE2-Luc) in MEFs treated with TGF-β1 and EHNA and grown on 0.5 kPa PAGs. Data represent the mean of two independent experiments ± S.E.M. \*\**p* < 0.01, \*\*\**p* < 0.005; n.s. indicates non-significant. E RT-qPCR analysis of selected genes using the same conditions as those shown in panel (A). Bar graph shows log_2_ fold change normalized to WT control. Data represent the mean of three independent experiments ± S.E.M. \**p* < 0.05, \*\*\**p* < 0.005; n.s. indicates non-significant as compared with WT (control). ^a^*p* < 0.05, ^b^*p* < 0.005; # indicates non-significant as compared with WT treated with TGF-β1. F WT MEFs were serum starved for 12 h and seeded into the collagen matrix. After 1 h, media containing TGF-β1 and/or EHNA were added into the matrices. Data represent the mean of two independent experiments ± S.E.M. \**p* < 0.05, \*\**p* < 0.01, \*\*\**p* < 0.005.

Next, we tested whether inhibition of dynein activity influences TGF-β1-induced transcriptional activity of YAP and Smad2/3. Treatment with EHNA dramatically suppressed the transcriptional activity of YAP/TEAD and Smad2/3, which was confirmed using reporter gene activity (Fig 6D); however, treatment with EHNA did not change the amount of acetylated microtubules (Figure EV6). Furthermore, the expression of myofibroblast marker genes, which are upregulated by TGF-β1 in MEFs grown on a soft substrate, was significantly decreased by treatment with EHNA (Fig 6E); therefore, 3D FMC was also greatly decreased (Fig 6F).

It has reported that downregulation of Lissencephaly-1 (Lis1) as a part of dynein complex abrogates the dynein function (Baumbach, Murthy et al., 2017). We also found that knockdown of Lis1 in WT MEF significantily decreased the TGF-β1 induced transcriptional activity of YAP and Smad and cellular contractile ability upon TGF-β1 stimulation (Figure EV7A-C). Altogether, these results indicate that dynein plays a critical role in acetylated microtubule-mediated nuclear translocation of YAP upon stimulation with TGF-β1, which, in turn, initiates the myofibroblast differentiation along with Samd2/3 activity in MEFs grown on soft substrates.

### Acetylated microtubules are critical for the progression of CCl_4_-induced hepatic fibrosis

Our abovementioned results indicate that acetylated microtubules initiate myofibroblast differentiation in MEFs grown on a soft substrate. Based on this information, we next used a CCl_4_-derived model of hepatic fibrosis to investigate whether acetylated microtubules appear in the early stage of fibrosis (Han, Koo et al., 2016). After mice were intraperitoneally (i.p.) injected with CCl_4_ for 2 weeks, the amount of acetylated microtubule was remarkably increased in the liver tissue of CCl_4_-treated mice compared with that in the livers of vehicle (corn oil)-treated mice; however, the expression of α-SMA was not significantly induced. As fibrosis progressed, the number of acetylated-α-tubulin positive cells was significantly increased, and these cells were dispersed in a radial arrangement (Fig 7A). Notably, this CCl_4_-induced increase in acetylated microtubules was abrogated by treatment with SB431542, a specific inhibitor of TGF-β receptor (TGFR) (Fig 7B). Western blotting, using samples shown in Fig 7B, confirmed that the presence of acetylated microtubules and α-SMA expression, upregulated by CCl_4_, were reduced in the mice treated with the TGFR inhibitor SB431542, suggesting that TGF-β1 is a potent agonist that promotes formation of acetylated microtubules in the early stage of liver fibrosis (Fig 7C).

**Figure 7.**
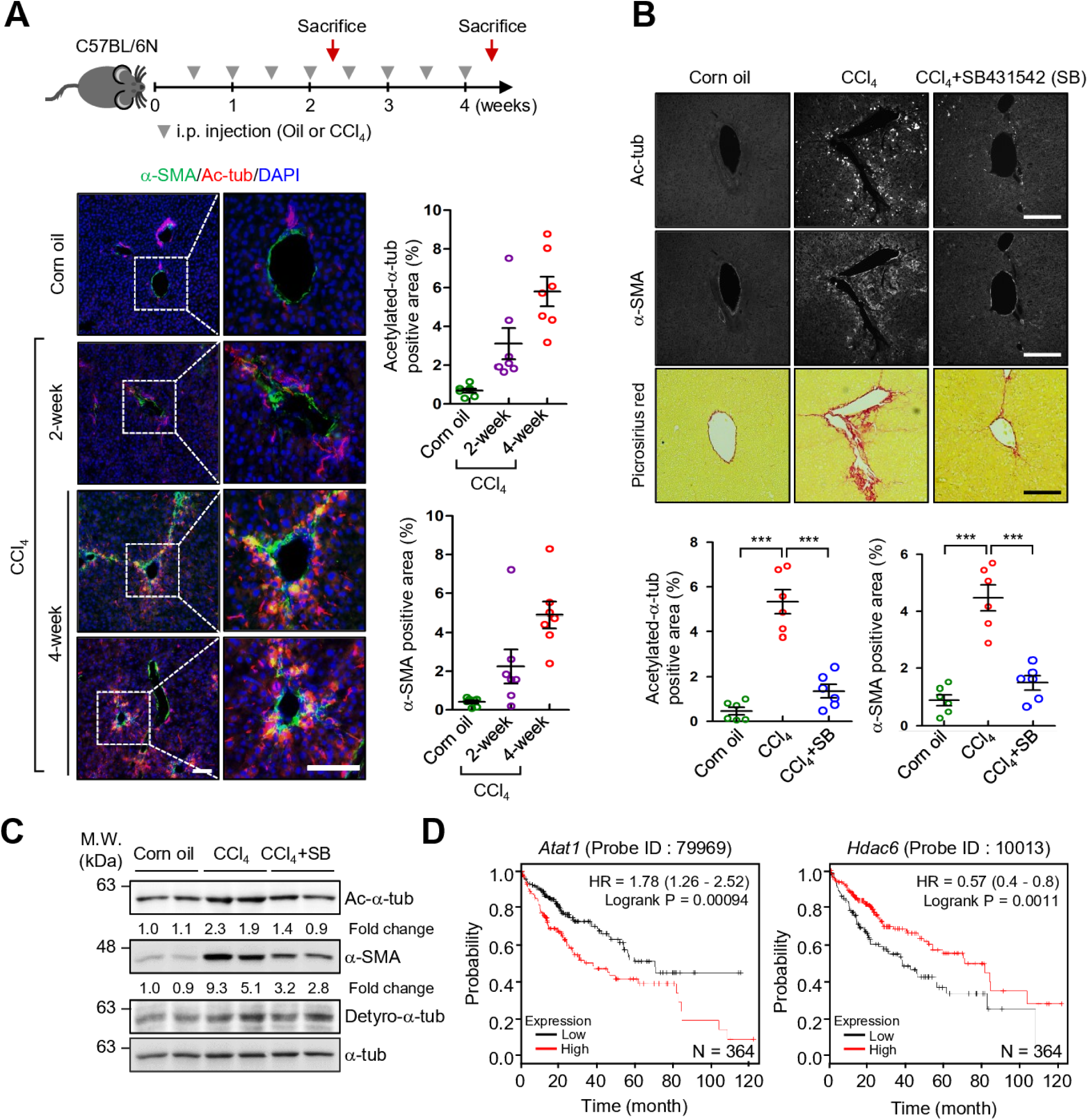
Acetylation of α-tubulin is initiated during CCl_4_-induced hepatic fibrosis. A Immunohistochemical (IHC) analysis of α-SMA and acetylated-α-tubulin expression in liver sections obtained from mice treated with corn oil or CCl_4_ (0.5 ml kg^-1^ body weight, i.p., twice a week) for 2 and 4 weeks. Graphs show the percentage of acetylated-α-tubulin and α-SMA positive area in each liver section. Scale bar, 200 μm. B IHC analysis of α-SMA and acetylated-α-tubulin expression, and picrosirius red stain for thick-collagen expression, in liver sections of mice administered combined treatment with SB431542 and CCl_4_ (10 mg kg^-1^ body weight, i.p., twice a week for 4 weeks) Representative images derived from replicate experiments (n = 3 each) were shown. Quantification of α-SMA and acetylated-α-tubulin-positive cells in liver tissue of mice treated as described above. Statistical significance of the differences between each treatment group and vehicle-treated group (\*\*\**p* < 0.005). Scale bar, 200 μm. C Western blotting for detection of acetylated-α-tubulin, detyrosinated-α-tubulin, and α-SMA in the liver sample obtained from (B). D Survival analysis was conducted using Kaplan-Meier survival curves with respect to the expression level of *Atat1* (probe ID; 79969) and *Hdac6* (probe ID; 10013) in patients with hepatocellular carcinoma (HCC) (n = 364).

We next aimed to elucidate the relation between a set of genes regulated by acetylated microtubules and the expression of *Atat1* or *Hdac6* (which encode enzymes that catalyse α-tubulin deacetylation) in 40 liver cirrhosis patients (GSE25097). A significant positive correlation between *Atat1* and the target genes including *Acta2, Tpm1, Postn, Loxl2, Cxcr6* was observed in the liver tissue (Figure EV8A-G). In contrast, the mRNA expression level of these genes showed a reverse correlation with that of *Hdac6* (Figure EV8A-G). Although *Ctgf* and *Cyr61* had high *p*-values, these genes also showed tendency of positive and reverse correlation with *Atat1* and *Hdac6* mRNA levels. The expression of *Yap1* was not correlated with those of *Atat1* and *Hdac6* (Figure EV8H).

Finally, we compared the survival rate in patients with fibrotic disease according to the expression level of *Atat1*. Although we were not able to analyse the survival rate in patients with liver fibrosis, we analyzed the survival rate in high-risk hepatocellular carcinoma (HCC) patients (n = 364, probe ID; 79989 and 10013) using Kaplan-Meier plotter. Patients with HCC also have liver cirrhosis, which develops after a long periods of chronic liver disease (Fattovich, Stroffolini et al., 2004). Our results indicate clinical relevance between high levels of acetylated-microtubule and low survival rate in patient with HCC (Fig 7D). Altogether, our findings demonstrate that TGF-β1 in an inflammatory environment induces fibrotic disease as a consequence of promotion of microtubule acetylation. Our results suggest that targeting of acetylated-α-tubulin can be a novel therapeutic approach to overcome fibrotic disease.

## Discussion

The increasing stiffness of normal stroma may provide a permissive environment for fibrotic disease. Myofibroblasts are the main cells that regulate the mechanical properties of tissue (Gabbiani, 2003, Hinz, 2007). Differentiation of fibroblasts into myofibroblasts has been studied extensively in association with fibrosis and wound healing. Studies examining myofibroblast differentiation *in vitro* indicate that myofibroblast differentiation requires a rigid ECM. However, one of the functions of myofibroblasts *in vivo* is to alter the stiffness of the ECM to render the tissue stiffer in pathological situations (Liu, Lagares et al., 2015); therefore, myofibroblast differentiation *in vivo* should initiate under conditions of soft ECM. Little is known about how myofibroblast differentiation occurs in an environment of mechanically soft ECM. In this study, we demonstrated that microtubule acetylation, induced by TGF-β1, is crucial for the expression of a set of myofibroblast marker genes. The expression of these genes, regulated via YAP and Smad2/3 activity in soft matrices, initiates myofibroblast differentiation.

The contractile force, exerted by cells, is increased by actomyosin activity after cells bind to ECM via activated integrin and focal adhesion structures (Chan, Chaudary et al., 2010). When cells interact with mechanically stiff substrates, the activity of actomyosin and focal adhesions is increased by the Rho signalling pathway via activated integrin, resulting in increased tension and contractility of the cells. Conversely, in a soft environment such as normal tissue, a cell shows minimal integrin-dependent actomyosin activity; consequently, the contractile force exerted by the cell is relatively weak (Rhee & Grinnell, 2007). In a soft matrix, cells are placed in non-tensional state; therefore, gene expression is minimal. For this reason, cells must acquire specialized structures that are responsible for cell differentiation under conditions of soft ECM. The results of this study suggest that acetylated microtubules participate in regulating gene expression and myofibroblast differentiation under condition of soft ECM. Our results, obtained using RNA-seq, reveal that acetylation of microtubules, induced by TGF-β1 under soft matrix conditions, increased the expression of approximately 83 genes that are involved in cytoskeletal reorganization and cellular processes. Among the genes, *Acta2*, *Tpm1* and *Tagln* are involved in cellular contractility (Gimona et al., 1992, Hinz, Celetta et al., 2001, Schevzov, Lloyd et al., 1993). We also found that the expression of *Loxl2*, encoding lysyl oxidase like-2 (LOXL2), which promotes ECM crosslinking and stabilization of the fibrotic matrix (Wong, Tse et al., 2014), is increased by acetylation of microtubules in soft matrices. Indeed, the *Loxl2* gene is significantly upregulated in fibrotic liver tissues (Wong et al., 2014). Based on this evidence, it is possible that structural remodelling of ECM via upregulation of these gene by acetylated microtubule in early fibrotic tissues may generate a positive feedback loop. This loop would further induce the myofibroblast activation accompanied by intrinsic mechanotransduction signalling of YAP and myocardin-related transcription factor/serum response factor (MRTF/SRF) via Rho/ROCK-dependent cytoskeletal reorganization (Calvo et al., 2013, Esnault, Stewart et al., 2014, Liu et al., 2015). The expression of some of these genes, increased by acetylation of microtubules upon stimulation with TGF-β1, plays an important role in transforming the mechanical environment of soft tissue into a stiff environment during the process of fibrosis. Hence, acetylation of microtubules, induced by TGF-β1 in the soft environment, is an essential factor for controlling the mechanical properties of tissues.

Recent reports have shown that microtubule acetylation in MEFs is transiently increased by stimulation with TGF-β1. In MEFs and COS-7 cells, the carboxyl region of α-TAT1 acts as a regulatory domain in which TGF-β1-associated kinase1 (TAK1) can directly bind to, and phosphorylates, the Ser237 residue upon stimulation with TGF-β1; therefore, acetylation of microtubules is increased upon stimulation with TGF-β1 (Shah, Kumar et al., 2018). However, we were unable to induce microtubule acetylation with stimulation by TGF-β1 under stiff 2D conditions (Fig 1D). This phenomenon has also been confirmed by other studies, showing that the amount of acetylated microtubules in cells such as fibroblasts, which originate from mesenchymal cells, is always upregulated regardless of the presence of growth factors when cells are under 2D stiff conditions (Gu, Liu et al., 2016). Our results obviously show that the amount of acetylated microtubules is dramatically increased under soft matrix conditions and in the presence of growth factors involved in myofibroblast differentiation such as LPA or TGF-β1. According to the four-quadrant cell-matrix system (Rhee & Grinnell, 2007), TGF-β1 signalling in soft ECM likely behaves differently than it does under stiff ECM conditions. TGF-β binds to the TGF-β receptor (TGFR), which possesses serine/threonine kinase activity and controls the various intracellular signalling pathways (Derynck & Zhang, 2003). As reported previously, catalytic activity of α-TAT1 is controlled by phosphorylation of its relatively long c-terminal regulatory domain. Therefore, examining how downstream kinases of TGFR, such as extracellular signal-regulated kinase, p38 mitogen-activated protein kinase, and casein kinases (Kim & Hwan Kim, 2013) regulate α-TAT1 activity in soft substrate condition will help to develop new treatments against fibrotic disease.

In this study, we have shown that the nuclear translocation of YAP in response to stimulation with TGF-β1 is required for the formation of acetylated microtubule/dynein complex in soft substrates. Recent studies have reported that microtubule and actin cytoskeleton networks are required for the nuclear translocation of numerous virus-and cancer-related proteins (Bremner, Scherer et al., 2009, Giustiniani, Daire et al., 2009). Microtubule binding protein p53 and PTH-related peptide (PTHrP) are translocated into the nucleus via interaction with importin β1 and importin α/β respectively (Lam, Briggs et al., 1999, Roth, Moseley et al., 2007). Heat shock protein 90 (Hsp90) also requires acetylated microtubule-based transport to enter the nucleus (Giustiniani et al., 2009). Fibroblast activation in the soft matrix is closely related to the characteristics of the microtubule network. Nuclear translocation of proteins in a microtubule-dependent manner is considered a major mechanism for the nuclear transport of proteins in a soft substrate. Interestingly, we also found that inhibition of dynein function by a pharmacological inhibitor or overexpression of dynamitin significantly reduced the nuclear translocation of the YAP protein, indicating that dynein plays a role in the nuclear translocation of proteins. Some NLS-containing proteins can bind the dynein motor complex via interactions with dynein light chains. The NLS-containing protein then utilises the dynein motor complex to travel along the microtubules towards the nucleus (Wagstaff & Jans, 2009). Although YAP does not contain a canonical NLS, the N-terminal 1-55 amino acids of Yorkie (YAP homology in *Drosophila*), especially Arg-15, are essential for the nuclear localization of YAP via direct interaction with importin-α1 (Wang, Lu et al., 2016). It is plausible that the interaction of YAP with microtubule/dynein complex in soft substrates leads to an accumulation of YAP in the nuclear periphery, where YAP containing non-canonical NLS is recognized by importin proteins.

Nevertheless, it is not clear how acetylated microtubules directly participate in the nuclear translocation of YAP in soft matrices. Recent studies have shown that the nuclear flattening, induced by ECM mechanical forces, is enough to initiate the nuclear translocation of YAP (Elosegui-Artola, Andreu et al., 2017). In a stiff matrix, YAP is translocated into the nucleus by ECM-nuclear mechanical coupling via linker of nucleoskeleton and cytoskeleton (LINC) complex. In our study, we observed changes in nuclear shape of MEFs stimulated with TGF-β1 under soft matrix conditions; these nuclear changes were inhibited by depletion of α-TAT1. In cells grown on a soft matrix, the nucleus is mechanically uncoupled from the ECM and is not exposed to external forces. In cells grown on a soft matrix and stimulated with TGF-β1, the increased tubulin acetylation results in altered nuclear shape, generating elongation (Fig 1A and Fig 2B). It is possible that tubulin acetylation provides mechanical support and dynein-dependent transport to cells stimulated with TGF-β1; this may constitute a mechanism for regulating YAP nuclear translocation after stimulation with TGF-β1 under soft matrix conditions. Therefore, our next challenge is to assess changes in the nuclear pore induced by acetylated microtubule in cells stimulated with TGF-β1.

The development of liver fibrosis is accompanied by progressive changes in the liver microenvironment. Liver fibrosis is strongly associated with excessive accumulation of ECM proteins, including collagen, resulting in cirrhosis, portal hypertension, and liver failure severe enough to require liver transplantation (Bataller & Brenner, 2005). The administration of CCl_4_ or bile duct ligation causes liver damage and induces the expression of factors such as TGF-β1, resulting in activation of hepatic stellate cells (HSCs) and fibroblasts induced via Smad phosphorylation (Han et al., 2016, Yin, Evason et al., 2013). Additionally, the nuclear localization of YAP is found in the HSCs of mice with CCl_4_-induced liver fibrosis. Inhibition of YAP signalling with verteporfin suppresses liver fibrosis and HSC activation via decreased expression of marker genes, such as *Acta2* and *Ctgf*, involved in HSC activation (Mannaerts, Leite et al., 2015). In this study, when C57BL/6N mice were injected with CCl_4_ for 2 weeks, the number of acetylated microtubule-positive cells in liver tissue were increased around portal vein and showed a radial shape, although α-SMA expression was not yet detectible. After CCl_4_ was administrated for 4 weeks, the fibrotic region positive for α-SMA expression was drastically expanded and the number of cells positive for acetylated microtubules had increased. The increase in tubulin acetylation during early fibrosis mediates the expression of myofibroblast marker genes and is associated with changes in ECM composition and architecture, resulting in tissue stiffening. The results of our present study indicate that tubulin acetylation is a molecular marker for myofibroblast-derived pathogenesis in early fibrosis. Further, targeting of microtubule acetylation may be an effective strategy for the treatment of pathogenic conditions such as fibrosis.

## Materials and Methods

### Antibodies and reagents

In this study, we used antibodies against α-SMA (Sigma-Aldrich; #A5228), acetylated-α-tubulin (Cell Signaling Tech., Beverly, MA, USA; #5335), detyrosinated-α-tubulin (Millipore, Billerica, MA, USA, #MAB3201), α-tubulin (Sigma-Aldrich, #T9026), GFP (Santa Cruz Biotech.; #sc-9996), pMLC (Ser 19; Abcam, Cambridge, MA, USA, #ab64162), YAP (Santa Cruz Biotech., #sc-101199), phospho-Smad2 (Ser465/467)/Smad3 (Ser423/425) (Cell Signaling Tech.; #8828), Smad2/3 (Santa Cruz Biotech., #sc-133098) and dynein (Millipore, Billerica, MA, USA, #MAB1618). TGF-β1 (R&D systems, MN, USA; #240B), as well as EHNA (Tocris Bioscience, UK, #1261) and nocodazole (Sigma-Aldrich, Taufkirchen, Germany, #M1404), were used as growth factor or inhibitors.

### Animal models and procedures

Male C57BL/6N mice, 6 weeks of age, were purchased from DBL (Chungcheongbuk-do, Korea). Mice were injected with vehicle (corn oil) or CCl_4_ (0.5 ml kg^-1^ body weight, i.p.) twice a week for 2 or 4 weeks. SB431542 (10 mg kg^-1^ body weight, i.p.) was injected into mice twice a week for 4 weeks, 1 day before CCl_4_ injection. Mice were sacrificed 2 days after the final CCl_4_ injection. The Chung-Ang University Institutional Review Board (IRB) approved all procedures involving mice used in this study.

### Immunochemistry

Cells were seeded on fibronectin-coated 0.5 kPa PAG or glass 12-mm coverslips and incubated with or without TGF-β1 for 8 h. Samples were fixed with 3.7% paraformaldehyde for 15 min and permeabilized with 0.5% Triton X-100 in phosphate buffered saline (PBS) for 10 min. To block non-specific signals, coverslips were blocked with 2% bovine serum albumin (BSA) in PBST (0.1% of Triton X-100 in PBS) for 1 h. Cells were then incubated with indicated primary antibodies (1:50∼100) for 1 h, followed by incubation with the appropriate secondary antibodies for 1 h at room temperature (RT). Coverslips were mounted on glass slides with Fluoromount-G (Southern Biotechnology Associates, Birmingham, AL, USA) and analyzed using an Eclipse 80i fluorescence microscope (Nikon, Tokyo, Japan). Images were acquired using a digital camera (digital sightDS-Qi1Mc, Nikon) and NIS-Elements image analysis software (Nikon). Image processing was carried out using Photoshop 11.0 (Adobe Systems, San Jose, CA, USA).

### Immunohistochemistry

Liver tissues were fixed with 4% paraformaldehyde and cryo-protected in increasing concentrations of sucrose solution (10, 20, and 30%) until tissue sinks. The tissues were then embedded in O.C.T compound (Tissue-Tek) and sectioned at 15 μm. The sections were blocked with M.O.M.^TM^ blocking solution and normal goat serum (Vector Biolaboratories, Burlingame, CA, USA) for 1 h at RT. Tissues were then incubated with antibodies specific for α-SMA and Ac-α-tubulin. After removing the unbound primary antibody with PBS, the tissues were incubated with a secondary antibody in 2% normal goat serum at RT for 1 h. Afterwards, the tissues were washed in PBS and mounted with Fluoromount (Southern Biotechnology Associates, Birmingham, AL, USA). Fluorescence-positive area was analyzed using imaging software NIS-Elements advanced research (Nikon).

### Lentiviral production and infection

For virus production, HEK293T cells were co-transfected using lentiviral packaging plasmids (psPAX2 and pMD2.G) with a PLKO.1-blast plasmid (Addgene, Cambridge, MA, USA, #26655) containing shSmad2 or shYAP. After 24 h, the media were replaced with fresh media, and HEK293T were incubated for an additional 48 h. The media were collected and filtered using a 0.45 μm syringe filter. MEFs were treated with shRNA containing lentiviral particles with polybrene (final concentration 8 μg ml^-1^) to enhance viral transduction. After 48 h, cells were selected using 5 μg ml^-1^ blasticidin (Sigma-Aldrich, #15205) in growth medium. The oligonucleotide pairs used are listed as follows in Table EV1.

### Western blotting

Cells were lysed with lysis buffer containing 1% Nonidet P-40 (NP-40), 1% sodium dodecyl sulfate (SDS), 150 mM NaCl, 6 mM Na_2_HPO_4_, 4 mM NaH_2_PO_4_, 2 mM ethylenediaminetetraacetic acid (EDTA), 50 mM NaF, 1 mM Na_3_VO_4_, 1 mM 1,4-dithiothreitol (DTT), and 1 mM phenylmethylsulfonyl fluoride (PMSF). The membranes with bound proteins were incubated with the indicated primary antibodies for 12∼16 h at 4°C. Then, membranes were further incubated with horseradish peroxidase (HRP)-conjugated secondary antibodies (Jackson ImmunoResearch Laboratories, West Grove, PA, USA) for 1∼2 h at RT. Signals were developed using enhanced chemiluminescence (Bio-Rad, Hercules, CA, USA) reagents, and band density was measured using a Quantity One^®^ system (Bio-Rad).

### Generation of α-TAT1 KO cell line using CRISPR/Cas9

A 20-bp guide RNA sequence (5′-catggagttcccgttcgatg-3′), targeting genomic DNA within exon 1 of *Atat1*, was selected from a Genescript database of predicted high-specificity protospacer adjacent motif (PAM) sequence in the mouse exome. PX458 or PX458 containing gRNA and PLKO.1-puro (Addgene, #10878), were co-transfected into MEFs. Transfected cells were selected with puromycin (2 μg ml^-1^) for 2 weeks, and single-cell colonies were acquired. α-TAT1 knockout in MEFs was verified by genomic DNA sequencing and western blotting.

### Luciferase reporter assay

MEFs were transfected with 4 μg of 8xGTIIC-luciferase (Addgene, #34615), SBE2-luciferase (Addgene, #16500), and 4 μg of pCMV-β-galactosidase (Clontech Laboratories, Inc., CA, USA) using electroporation according to the manufacturer’s protocol. After 24 h, the transfected cells were serum-starved for 12 h, and seeded on fibronectin-coated 0.5 kPa PAGs for 8 h with or without TGF-β1 (2 ng ml^-1^). Cells were lysed using reporter lysis buffer (Promega, WI, USA), and lysates were analyzed using a GloMax^®^ Luminometer (Promega). Transfection and expression efficiencies were normalized to the activity of β-galactosidase activities.

### Quantitative real-time PCR

Total RNA was extracted using RNAiso Plus reagent (TaKaRa, Tokyo, Japan; #9109), and complementary DNA was synthesized using M-MLV reverse transcriptase (M. Biotech, Seoul, Korea). qPCR was conducted with SYBR Premix Ex-Taq II (Tli RNase H Plus, TaKaRa, #RR820A) per manufacturer’s instructions on a Quantstudio^TM^3 instrument (Applied Biosystems) using primers listed in Table EV2. The expression level of each gene was calculated as 2^−*ΔΔCt*^ and normalized to the Ct value of *Gapdh*. The results were obtained using three biological replicates and two or three technical replicates for each gene and sample.

### Library preparation and RNA-sequencing

After total RNA was extracted from WT and α-TAT1 KO grown on soft matrix with or without TGF-β1, RNA quality was measured on an Agilent 2100 Bioanalyzer using an RNA 6000 Nano Chip (Agilent Technologies, Amstelveen, Netherlands). RNA quantification was performed using a ND-2000 Spectrophotometer. For control and test RNA, the construction of a library was performed using QuantSeq 3’ mRNA-Seq Library Prep Kit (Lexogen, Inc., Austria) according to the manufacturer’s instructions. Each 500-ng of total RNA was prepared, then an oligo-dT primer containing an Illumina-compatible sequence at its 5’ end was hybridized to the RNA, and reverse transcription was conducted. After degradation of the RNA template, second strand synthesis was initiated using a random primer containing an Illumina-compatible linker sequence at its 5’ end. The double-stranded library was purified using magnetic beads to remove all reaction components. The library was amplified to add the complete adapter sequences required for cluster generation. The finished library was purified from PCR components. High-throughput sequencing was performed as single-end 75 sequencing using NextSeq 500 (Illumina, Inc., USA).

### Bioinformatic analysis

QuantSeq 3’ mRNA-Seq reads were aligned using Bowtie2. Bowtie2 indices were either generated from the genome assembly sequence or the representative transcript sequences for aligning to the genome and transcriptome. The alignment file was used for assembling transcripts, estimating their abundance and detecting differential expression of genes. Differentially expressed genes were determined based on counts from unique and multiple alignments using coverage in Bedtools. The Read Count data were processed based on the Global normalization method using the Genowiz™ version 4.0.5.6 (Ocimum Biosolutions, India). Hierarchical clustering analysis was performed using Pearson correlation as the distance metric. Functional classification of gene on searches done by PANTHER gene ontology (http://www.pantherdb.org). Survival rate in HCC patients were analyzed by Kaplan-Meier plot (http://kmplot.com).

### Statistical analysis

The differences between control and treatment groups were statistically analyzed by Student’s *t*-test. Data are expressed as the mean ± standard error of the mean (S.E.M.) of three independent experiments; *p*-values less than 0.05 were considered statistically significant.

## Acknowledgement

This research was supported by the National Research Foundation of Korea (to Rhee, NRF-2017R1A2B4004324) and the Korea Healthcare Technology R&D Project, Ministry for Health & Welfare Affairs (to Rhee, HI17C1620).

## Author Contributions

E.Y. and S.R. designed the project and wrote the manuscript. E.Y. conducted the experiments. E.Y. and S.R. analyzed the results. J.J. and S.K. contributed to the experimental work. P.K. and Y.S. contributed to the mouse experimental work. J.-W.K. provided advice in the bioinformatic data. W.-K.S. and S.R. supervised and administered the project, and all authors critically revised the manuscript and approved its final version.

## Competing Interests

The authors declare that they have no competing interests.

## Expanded Table

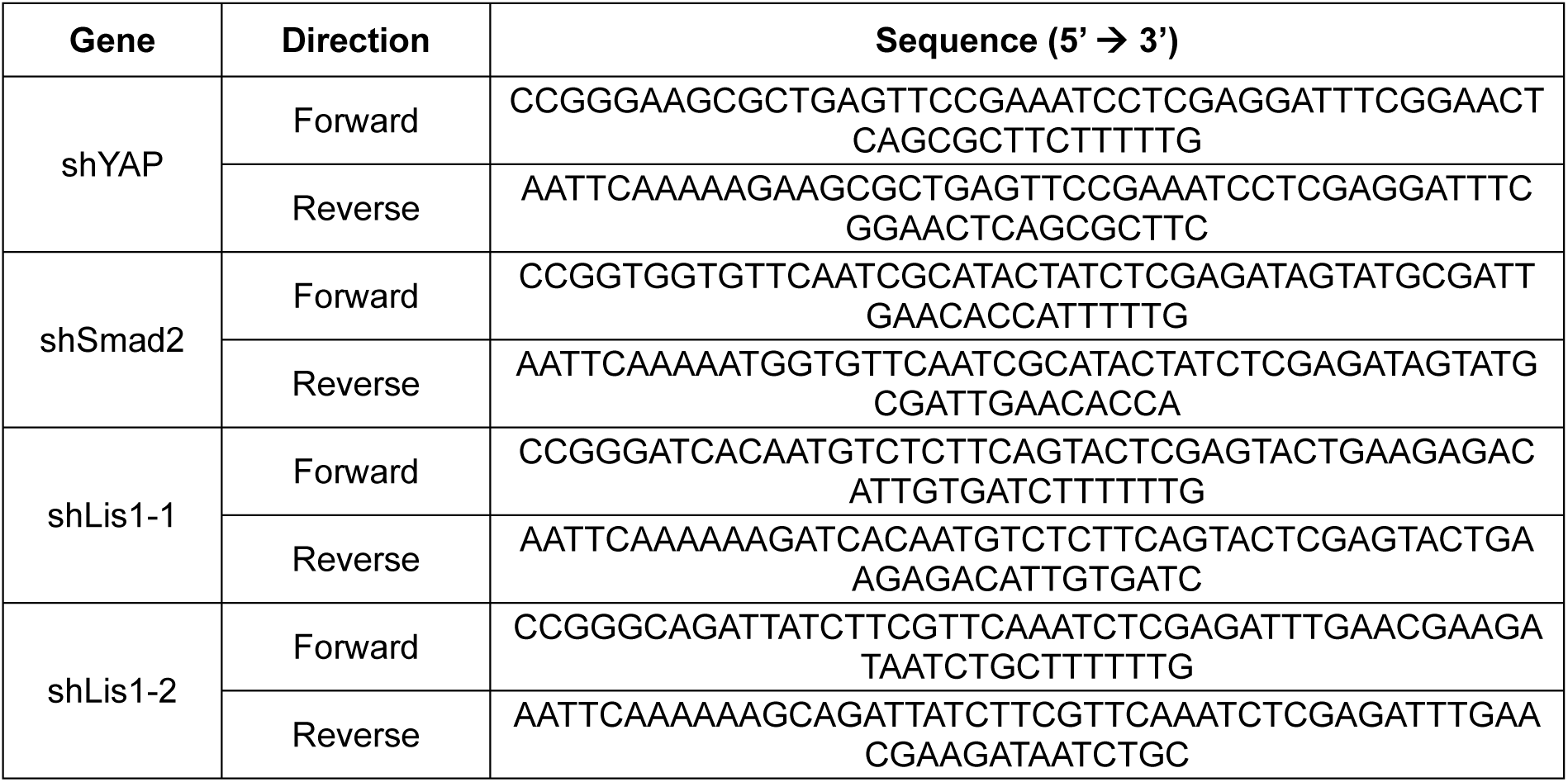
Table EV 1.

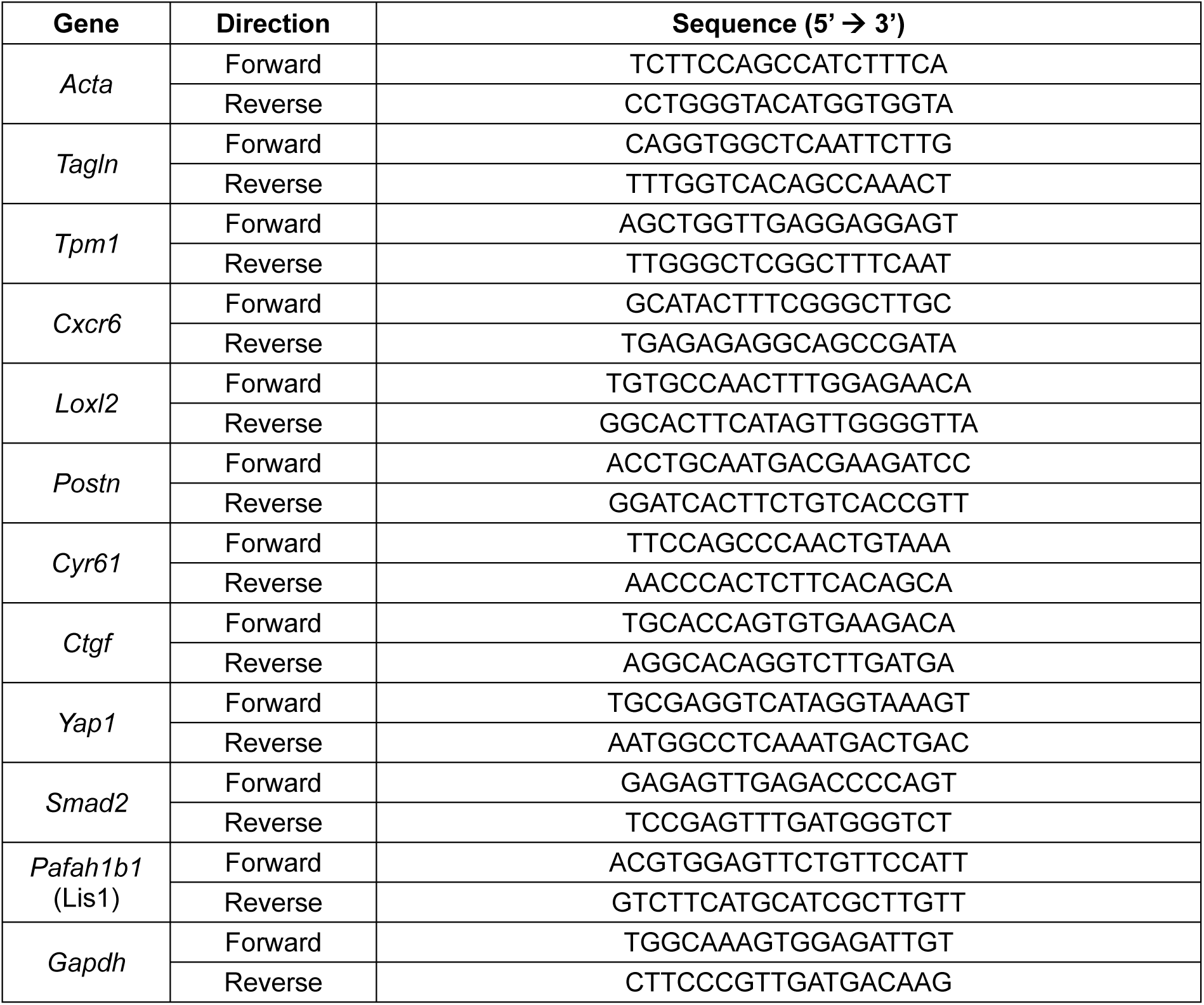
Table EV 2.

## Expanded Figure Legends

**Figure EV1. Increase in acetylated-α-tubulin in MEFs grown on soft matrices is correlated with α-SMA expression.**

A, B Cells were incubated on 0.5 kPa or glass and treated with TGF-β1 (2 ng ml^-1^), LPA (10 μM), or PDGF (10 ng ml^-1^) for 8 h. Then, western blotting was performed using antibodies specific for acetylated-α-tubulin and α-SMA.

**Figure EV2. Acetylated-α-tubulin is increased in low-tension environment.**

A Immunofluorescence imaging of MEFs, incubated on fibronectin-coated coverslips and treated with TGF-β1 combined with blebbistatin and Y27632 (1, 5, 10 μM), and immunolabeled with antibodies specific for acetylated-α-tubulin and vinculin. Arrow indicates long extended acetylated-α-tubulin. Scale bar, 50 μm.

B Western blotting for acetylated-α-tubulin and α-SMA in cells incubated as described in (A). Graph shows relative expression of acetylated-α-tubulin and α-SMA normalized to that of α-tubulin.

**Figure EV3. Generation of α-TAT1 KO MEFs, and comparison of α-SMA expression between WT and α-TAT KO.**

A Empty PX458 and guide RNA (gRNA) constructs used for insertion were transfected with B16F10 cells, and then selected with puromycin for 1 week. Genomic DNA obtained from puromycin resistant B16F10 cells was amplified (870 bp). PCR products were denatured, re-annealed, and incubated with T7 endonuclease (T7E1) to cleave mismatched DNA. PCR products were excised into two fragments (593 and 277 bp).

B Validation of α-TAT1 KO MEFs by genomic DNA sequencing.

C TGF-β1-induced *Acta2* (encoding α-SMA) expression was compared between WT and α-TAT1 KO MEFs in transcripts levels expressed under soft and stiff conditions. The graph represents the mean of three independent experiments ± S.E.M. \**p* < 0.05, \*\**p* < 0.01, \*\*\**p* < 0.005; n.s. indicates non-significant.

**Figure EV4. Microtubule acetylation increases YAP nuclear translocation in MEFs grown on a soft matrix.**

WT and α-TAT1 KO MEFs were seeded on fibronectin-coated soft and stiff matrices and stimulated with TGF-β1 for 8 h. Cells were fixed and labelled with antibodies specific for YAP and acetylated α-tubulin. Graphs show the quantification of YAP cellular localization in MEFs grown on soft and stiff matrices. N; nucleus, C; cytosol. Scale bar, 50 μm.

**Figure EV5. Overexpression of dynamitin suppresses TGF-β1-induced YAP nuclear translocation on soft matrices.**

A WT MEFs were transfected with GFP and GFP-dynamitin for 24 h and seeded on fibronectin-coated 0.5 kPa PAG and glass with or without TGF-β1 for 8 h. Cells were then fixed and immunocytochemistry was performed using an antibody specific for YAP. Dotted lines indicate GFP-expressing cells. Scale bar, 50 μm.

B Quantification of YAP cellular localization in MEFs grown on soft matrices. N; nucleus, C; cytosol.

**Figure EV6. Inhibition of dynein activity does not affect TGF-β1-induced microtubule acetylation and total Smad phosphorylation in MEFs grown on soft matrices.**

MEFs were incubated on a soft matrix and treated with TGF-β1 and/or EHNA for 8 h. Then the cells were lysed and subjected to western blotting using antibodies specific for α-SMA, acetylated α-tubulin, and phospho-Smad2/3.

**Figure EV7. Knockdown of Lis1 to inhibit the dynein function suppresses transcriptional activities of YAP and Smad upon TGF-β1 on soft matrices.**

A Generation of Lis1 knockdown cell line using shRNA lentiviral infection. Knockdown efficiency of Lis1 was verified by RT-qPCR. \*\*\**p* < 0.005, n.s. non-significant.

B Transcriptional activity of YAP and Smad was compared by luciferase assay in Lis1 KD cell lines upon TGF-β1 stimulation on soft matrix. \*\**p* < 0.01, \*\*\**p* < 0.005, n.s. non-significant.

C Collagen matrices containing Mock and Lis1 KD cells (shLis1-1 and shLis1-2) were incubated for 12 h under floating condition with TGF-β1. Contractility of cells was measured the reduced perimeter of collagen gel. \*\**p* < 0.01, \*\*\**p* < 0.005, n.s. non-significant.

**Figure EV8. Correlation plot of genes expression regulated by α-TAT1 depending on *Atat1* and *Hdac6* in liver fibrosis tissue.**

A-H mRNA expression level of *Acta2, Tpm1, Loxl2, Cxcr6, Postn, Cyr61, Ctgf* and *Yap1* with respect to *Atat1* and *Hdac6* mRNA expression in the liver tissue (40 liver cirrhosis patients; GSE25097). Pearson correlation coefficient (r) and *p*-value (*p*) from r were calculated.

